# Detecting seabird responses to invasive species eradication

**DOI:** 10.1101/2021.04.07.438876

**Authors:** Jeremy P. Bird, Richard A. Fuller, Aleks Terauds, Julie C. McInnes, Penelope P. Pascoe, Toby D. Travers, Rachael Alderman, Justine D. Shaw

## Abstract

Maximising survey efficiency can help reduce the trade-off between spending limited conservation resources on evaluating performance of past interventions and directing those resources towards future interventions. Seabird responses to island eradications are often poorly evaluated owing to financial, logistical and methodological challenges associated with remote field work and species ecology. We surveyed an assemblage of threatened seabirds following the world’s largest island eradication of multiple invasive species, testing multiple survey designs and outputs. We compared the outcomes of two important choices made during survey design: 1) whether to use unbiased or targeted surveys; and 2) implementing design-based or model-based analyses. An unbiased whole-island stratified randomised survey design performed well in terms of confidence in the final population estimates for widespread species, but poorly for localised recolonising species. For widespread species, model-based analyses resulted in slightly lower population estimates with narrower confidence intervals than traditional design-based approaches but failed to capture the realised niches of recolonising species, resulting in population estimates three orders of magnitude higher than current best estimates. We conclude that a multi-method approach to survey design best captures the size and distribution of recovering populations when the study system is ecologically diverse—importantly our results suggest there is no single strategy for efficient surveys of diverse seabird communities following large island invasive species eradications.

## INTRODUCTION

The removal of invasive species from islands has been used effectively to prevent species extinctions and promote recovery of native wildlife (Courchamp et al. 2014, Jones et al. 2016, Bolam et al. 2020). Post-eradication biodiversity assessments are needed to inform funders, policy-makers and the public of outcomes, and to help plan and prioritize future eradications by understanding ecological responses (Towns 2018). However, funding for eradication efforts focuses on the removal of invasive species, with limited resources allocated for post-eradication monitoring (e.g. Springer, 2016), and such funding is often available only in the years immediately following eradication. In this context, maximising survey efficiency to deliver the most information within financial and practical constraints is necessary (Possingham et al. 2012). Currently there are some key challenges to overcome when trying to detect post-eradication responses. First, recovery takes time, so population responses (and associated effect sizes) are typically small immediately following eradication, particularly amongst species that are slow to reproduce (Brooke et al. 2018). Second, the size, scale and complexity of eradication operations is increasing (Veitch et al. 2019) and detecting preliminary responses within large landscapes is challenging, especially among rare and recolonising species. Furthermore, recolonization events can occur stochastically (Cowie and Holland 2006), making them difficult to predict in space or time. Third, the remoteness of many islands where eradications have been completed introduces high costs and logistical challenges (Rodríguez et al. 2019).

Field and analytical methods are constantly evolving to help overcome these challenges (Thompson 2013), but a side-effect of this progress is the daunting array of alternative methods available to practitioners. Here we studied an assemblage of burrowing seabirds following the world’s largest multi-species eradication of invasive vertebrates to provide methodological insights and guidance. Together our study species and system present all the hallmark challenges outlined above.

Macquarie Island (54°30’S, 158°57’E) is a remote oceanic island approximately 1,500 km south-east of Tasmania, Australia. Following decades of impacts introduced Wekas *Gallirallus australis*, Cats *Felis catus*, Rabbits *Oryctolagus cuniculus*, Black Rats Rattus rattus and House Mice *Mus musculus* have been removed through staged management and eradication (Copson and Whinam 2001, Robinson and Copson 2014, Springer 2016). The island is large and the terrain accessible only by foot, so surveys require multi-day field trips. Its status as a UNESCO World Heritage Site and Tasmanian Nature Reserve mean access permits are required, and the sub-Antarctic environment and island’s terrain add logistical difficulty. The four procellariiform petrels in our study are diverse, comprising widespread common species, and localised rare species that have only recently recolonised the island in response to invasive species removal (Brothers and Bone 2008). All four species nest in under-ground burrows distributed discontinuously across challenging terrain, the entrances to which can be obscured by vegetation. Birds only enter or leave burrows at night, and species are not present year-round (Warham 1990, 1996). Owing to these behaviours, petrels are widely acknowledged to be a difficult group to survey (Rayner et al. 2007, Rodríguez et al. 2019), and yet because petrels are *K*-selected species and respond more slowly to eradications than other seabirds (Brooke et al. 2018), substantial statistical rigour is required to detect short-term changes.

The ultimate aim of our research is to provide data to inform stakeholders of the outcomes of invasive species eradication from Macquarie Island, and to understand species’ ecological responses, but here our aim is to answer the initial question: “what survey and analytical tools do you need in your toolkit to study seabird responses to eradication?”

One fundamental choice when designing surveys is whether to use an unbiased or purposefully targeted survey design. Unbiased designs such as stratification or randomisation are recommended where the goal of surveys is to estimate populations and track changes over time (Buckland et al. 2015). However, for rare or highly localised species that can easily be missed within the landscape, they may result in low precision (Thompson 2013, Pacifici et al. 2016). In such cases using prior knowledge such as historical records, pilot ground surveys and nocturnal surveys of aerial activity can improve results (Rayner et al. 2007, Arneill et al. 2019).

Additionally, when analysing data, design-based or model-based analyses can be used. Model-based methods rely on the relationship between species abundance and spatial and environmental covariates to infer abundance in space and/or time. They can improve precision in animal abundance estimates by eliminating biologically impossible changes in density through smoothing (Camp et al. 2020). While design-based methods can require stratification of survey data to account for uneven sampling (possibly by design), model-based methods utilise data more efficiently through finer scale covariates. However, switching from design- to model-based methods requires a substitution of assumptions: from assuming sampling locations were chosen at random, to assuming the model accurately reflects a species’ realised niche (Camp et al. 2020). Information is currently limited as to how well this assumption performs for different types of species, e.g. common or rare, and clustered or dispersed species (Pacifici et al. 2016, Dilley et al. 2019).

Estimating burrowing petrel populations and trends requires two fundamental metrics: the number of burrows, and the proportion that are occupied (Bird et al., submitted A). We focus here on methods for counting and estimating the total number of burrows (see Bird et al., submitted B for approaches to estimating burrow occupancy on Macquarie Island). We conducted parallel whole-island stratified randomised plot and transect surveys for all burrowing petrels and compared these to targeted whole-island search surveys for two rare recolonising species. To test the performance of these approaches when surveying a diverse fauna on a large island post-eradication, we 1) compared the performance of unbiased and biased survey designs for common widespread versus rare localised species, and 2) compared traditional design-based analyses with model-based analyses.

## METHODS

We generated population estimates and distribution maps for evaluation and reporting through five key steps: i) a preliminary survey to gain prior knowledge of species distributions and encounter rates; ii) island stratification and randomised survey design; iii) implementation of unbiased island-wide plot and transect surveys, and targeted surveys; iv) design- and model-based analyses of burrow numbers; and v) breeding population estimation by adjusting burrow estimates by occupancy (Figure 1). Our study focussed on two petrel species which persisted alongside invasive vertebrates, Antarctic Prions *Pachyptila desolata* and White-headed Petrels *Pterodroma lessonii*, and two recolonising species that were extirpated from the main island in the 1900s, Blue Petrels *Halobaena caerulea* and Grey Petrels *Procellaria cinerea* (Brothers 1984).

**Figure 1:**
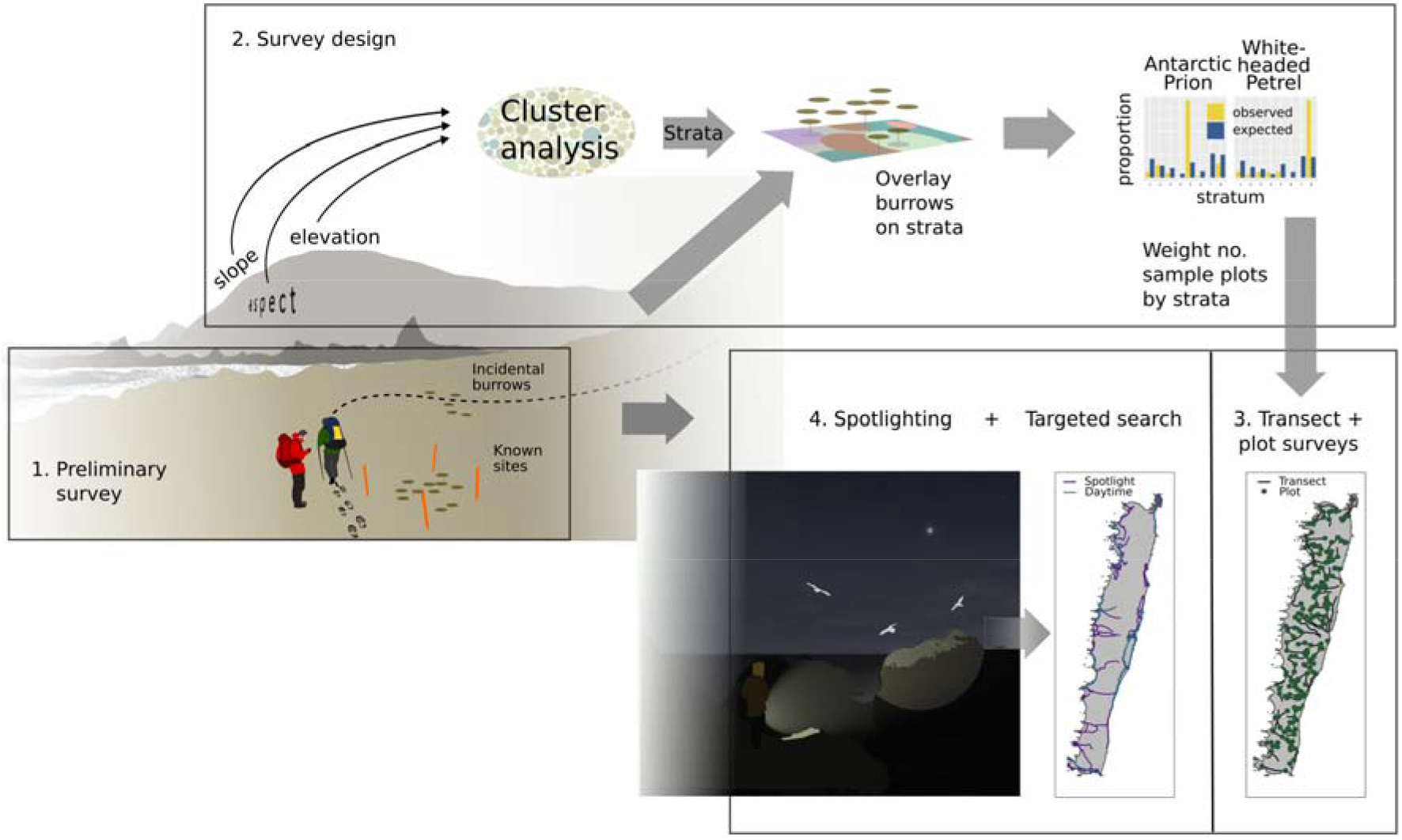
Petrel survey design and implementation on Macquarie Island. Burrows of different species were recorded at known sites and incidentally elsewhere (a). The frequency of incidental burrows in different modelled strata (b) was used to assign numbers of random plots in different strata, with transects walked between them (c). For localised species knowledge from existing sites was augmented through spotlight surveys to define daytime search areas (d).

### i) Preliminary survey

We undertook a preliminary survey in November-December 2017 to visit known petrel colonies and opportunistically record encounter rates in different habitats across Macquarie Island. All burrows encountered were identified to species level (see Bird et al. submitted B) and georeferenced using a handheld Garmin 60 GPS.

### ii) Environmental covariates, stratification and survey design

We stratified the island and used the distribution information collected during our preliminary survey to assign survey effort preferentially to strata with higher burrow encounter rates to minimise uncertainty around higher density estimates (see Supporting Information for full description of methods).

### iii) Survey implementation

#### Island-wide plot and transect surveys

From 5^th^ January to 24^th^ April 2018 we measured burrow density at 194 of the randomly generated points. At each sample location we thoroughly searched 100 m^2^ within a circle of radius 5.66 m (measured with a length of rope). All burrows were identified to species (see Bird et al. submitted B), counted and recorded by handheld GPS if the centre of the burrow entrance was within the plot.

To generate a second measure of burrow density we distance sampled burrows along 158 km of transects navigated between plots (see Supporting Information for full description of methods).

#### Targeted species-specific searches

For species not encountered incidentally during the preliminary survey, i.e. they were not widespread away from known colonies, we undertook targeted searches. In 2017 and 2018 we surveyed all known colonies of Blue Petrels and Grey Petrels (see Supporting Information for full description of methods). We attempted to census Grey Petrels, searching outwards from all located burrows to search the whole area of suitable habitat (Schulz et al. 2006). When surveying Blue Petrel colonies, we followed Dilley et al. (2017).

### iv) Analysis

#### Design-based analyses

##### Colony counts

For Blue and Grey Petrels we totalled all colony-level burrow estimates to generate a whole-island estimate.

#### Plot extrapolations

We digitised plots, clipped them by strata and calculated the area of each stratum surveyed within each plot. We assigned burrow waypoints to strata and calculated the density of burrows per stratum for each species in each plot. We multiplied the mean burrow density per stratum by the total area of each stratum to generate a total burrow estimate per stratum with upper and lower 95% confidence limits based upon bootstrapped percentiles (Derylo 2018). These were summed to give a whole-island burrow estimate with upper and lower limits.

#### Distance analyses

We manually cleaned transect tracks in BaseCamp by deleting extraneous points related to GPS scatter (when the observer was not moving). We imported the cleaned tracks into QGIS (QGIS Development Team 2019) and sub-divided them by vegetation height and strata. We then assigned all burrow observations to transect sections and recorded vegetation height and stratum as attributes. For Antarctic Prions and White-headed Petrels, we estimated average burrow density within each stratum along transects. We used detection probabilities to adjust raw burrow counts based upon the distance-based sampling assumption of uniform distribution within each transect section (Miller et al. 2017). We truncated the data, with the most distant 5% of observed burrows excluded from analyses. We then fitted half normal and hazard rate detection functions with cosine, hermite and polynomial adjustments and with stratum, observer and vegetation height as covariates. To account for overdispersion of our distance data we used a two-step process to select a final detection function using an adjust version of Akaike’s Information Criterion (QAIC) following Howe et al. (2019). We used the selected model to estimate abundance for each stratum within the area covered by transects (twice the truncation distance multiplied by transect lengths) and then scaled this to provide an overall burrow estimate and upper and lower confidence limits for the whole island (Miller et al. 2017).

#### Model-based analyses

We used a generalised additive model (GAM) and the detection probability functions described above to build density surface models (DSMs) for Antarctic Prions and White-headed Petrels from distance sampling data and environmental covariates (Miller et al. 2013). After selecting a detection probability function (stage 1), for stage 2, we clipped our transects into sections 2 × the truncation distance in length—5.5 m for analysis of Antarctic Prion data and 11.10 m for White-headed Petrel data. The sections were then buffered by the truncation distance to create contiguous roughly square sections of strip transect. Observations were joined to their respective transect sections to generate section-specific burrow counts. Mean values of the same environmental variables used to generate strata were calculated for each transect section as seven individual covariates. Generalized additive models (GAMs) were fitted in the ‘dsm’ package (Miller et al. 2020) with estimated abundance—burrow counts corrected for detection—as the response variable. We constructed global models with estimated abundance, spatial smooths of all seven environmental variables, and a bivariate spatial smooth of *x* and *y* coordinates from burrow observations (Marshall et al., 2017). Three global models were fitted using quasipoisson, negative binomial and tweedie distributions. The default basis complexity (*k = 10*) was used for all variables except the bivariate *xy* smooth which was assigned a basis complexity of *k = 100* to allow a high level of spatial complexity and account for the spatial autocorrelated distribution of burrows (Marshall et al. 2017). We selected a probability distribution based upon diagnostic plots and adjusted *r^2^* from the three global models and compared all subsets of the global model using AIC. The final model (lowest AIC score) was tested for concurvity (Marshall et al. 2017, Bock 2018). The final model for each species was used to predict burrow abundance across the entire island at a 20 × 20 m grid cell resolution. The density values in all cells were summed to produce an island-wide estimate of burrow numbers. We used the delta method for estimating 95% confidence intervals, and the variance propagation method when they were (Bravington et al. 2019, Miller et al. 2020).

We also generated spatial models of Blue and Grey Petrel abundance. Our data for these two species were derived from targeted surveys. Unlike in distance sampling where uncertain detection is explicitly accounted for, we assumed perfect detection of burrows during our targeted searches. Therefore, there was no need to incorporate detection probability. We extracted environmental covariates in the same way as above, and fitted GAMs directly. As above we used these to predict abundance island-wide.

All analyses were performed in ‘R’ using associated packages ‘e1071’, ‘distance’, ‘dsm’, ‘mgcv’, ‘tabularaster’ and ‘tidyverse’ (Wood 2017, Miller et al. 2017, 2020, Sumner 2018, Meyer et al. 2019, Wickham et al. 2019, R Core Team 2020).

### v) Population estimation

We adjusted burrow estimates by species-specific occupancy from Bird et al. (submitted B) to generate population estimates. Occupancy was not modelled spatially but was used as a constant to adjust the burrow estimates. We multiplied the estimated means to generate a point estimate of population size. We used the delta method to combine variances around the burrow and occupancy estimates before calculating confidence intervals.

### Data deposition

All data and R code are available from the Australian Antarctic Data Centre: <http://XXXXX>.

## RESULTS

### Preliminary survey and design

We recorded 1,521 burrows during our preliminary survey including known sites and incidental observations. Antarctic Prions and White-headed Petrels were the only widespread species with 696 and 467 burrows found respectively (Figure 1), so we optimised effort in the plot and transect surveys for the two widespread species. When overlain on our modelled strata Antarctic Prions and White-headed Petrels showed a strongly skewed distribution (Figure 1), which resulted in an increased survey effort on the plateau and upper escarpment slopes in the final design.

### Survey implementation

During the plot survey we counted 94 Antarctic Prion burrows in 25 of the 196 plots surveyed and eight White-headed Petrel burrows in five plots. No other species’ burrows were encountered. In the simultaneous survey of transects between each plot we recorded 2,845 Antarctic Prion burrows, 306 White-headed Petrel burrows and two Blue Petrel burrows. An additional 90 burrows (or just under 3%) were not identified to species-level so were excluded from analyses.

Having encountered just two Blue Petrel burrows and no Grey Petrel burrows during our island wide plot and transect surveys, we walked 71 km during nocturnal surveys and searched 249 km during follow-up ground searches targeting suitable habitat for these two species. During these searches we surveyed 37 Blue Petrel colonies in total including 12 newly identified sites, and 74 Grey Petrel colonies including 31 new sites.

### Detection probability estimation

A hazard rate key function with no adjustments and no covariates was selected by QAIC as the best detection function for Antarctic Prion burrows. The Kolmogorov-Smirnov goodness-of-fit test was not significant at the p = 0.05 level (*D_n_* = 0.03, p = 1) indicating a good fit between the model and the data. Truncation distance was w = 5.5 m. QAIC selected a half normal key function with observer as a covariate for White-headed Petrel burrows (Supporting Information). Again, the Kolmogorov-Smirnov goodness-of-fit test was not significant at the p = 0.05 level (*D_n_* = 0.04, p = 0.99).

### Survey coverage

The total area surveyed during our unbiased island-wide transect survey was 48 ha and 60 ha, or 0.4 and 0.5% of the whole island for Antarctic Prions and White-headed Petrels respectively (Figure 1). This comprised 2 × effective strip half width (ESW) × total transect length during transect surveys (Antarctic Prions: ESW = 3.04, SE = 0.07, 95% CI 2.90-3.18; White-headed Petrels: ESW = 3.82, SE = 0.21, 95% CI 3.68-3.96). The total areas searched for Blue and Grey Petrels were 1,192 and 1,264 ha, or 9.3 and 9.9% of the whole island comprising the island-wide transect surveys plus buffered ground searches and nocturnal search tracks.

### Design-based density estimation

Peak burrow densities recorded in plots were 0.14 Antarctic Prion burrows m^−2^, and 0.04 White-headed Petrel burrows m^−2^. Both the plot and transect data were used to extrapolate estimates of the total number of Antarctic Prion and White-headed Petrel burrows on the island (Table 1). Peak Blue Petrel burrow density recorded at a single colony was 1.79 burrows m^−2^ (Supporting Information). Grey Petrel numbers are still sufficiently low in identified colonies that we were able to count all burrows, resulting in a whole-island census of 630 burrows.

**Table 1:**
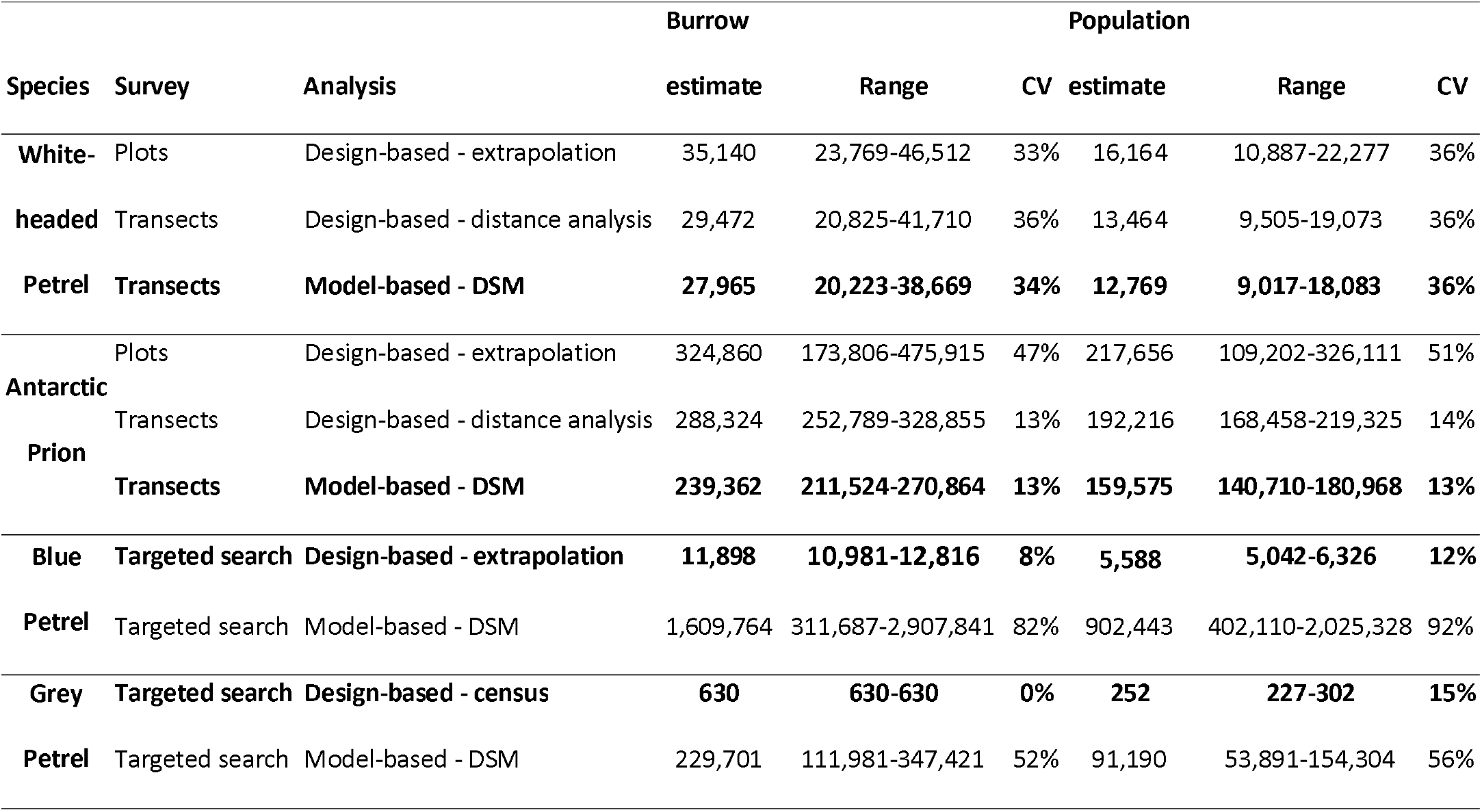
Estimates of the total number of burrows, breeding occupancy and total population estimates for four burrowing petrel species on Macquarie Island. The upper and lower confidence limits are expressed as a range, with the coefficient of variation for each estimate. The most precise estimates for each species are highlighted in bold.

### Model-based density estimation

The final DSM based on AIC for Antarctic prions included a bivariate smooth of xy co-ordinates, and smooths of elevation, NDVI and slope, with a negative binomial response distribution. The final DSM for White-headed Petrels included a bivariate smooth of xy co-ordinates, and smooths of elevation, NDVI, slope and wetness as predictors, with a Tweedie distribution (Supporting Information). Predicted burrow densities along transects peaked at 0.04 burrows m^−2^ for Antarctic Prions and 0.014 burrows m^−2^ for White-headed Petrels. DSMs were used to predict species abundance island-wide based upon the spatial environmental layers, resulting in a whole-island estimate of the number of burrows for comparison with estimates from design-based methods (Table 1) and generating species distribution maps (Figure 2).

**Figure 2:**
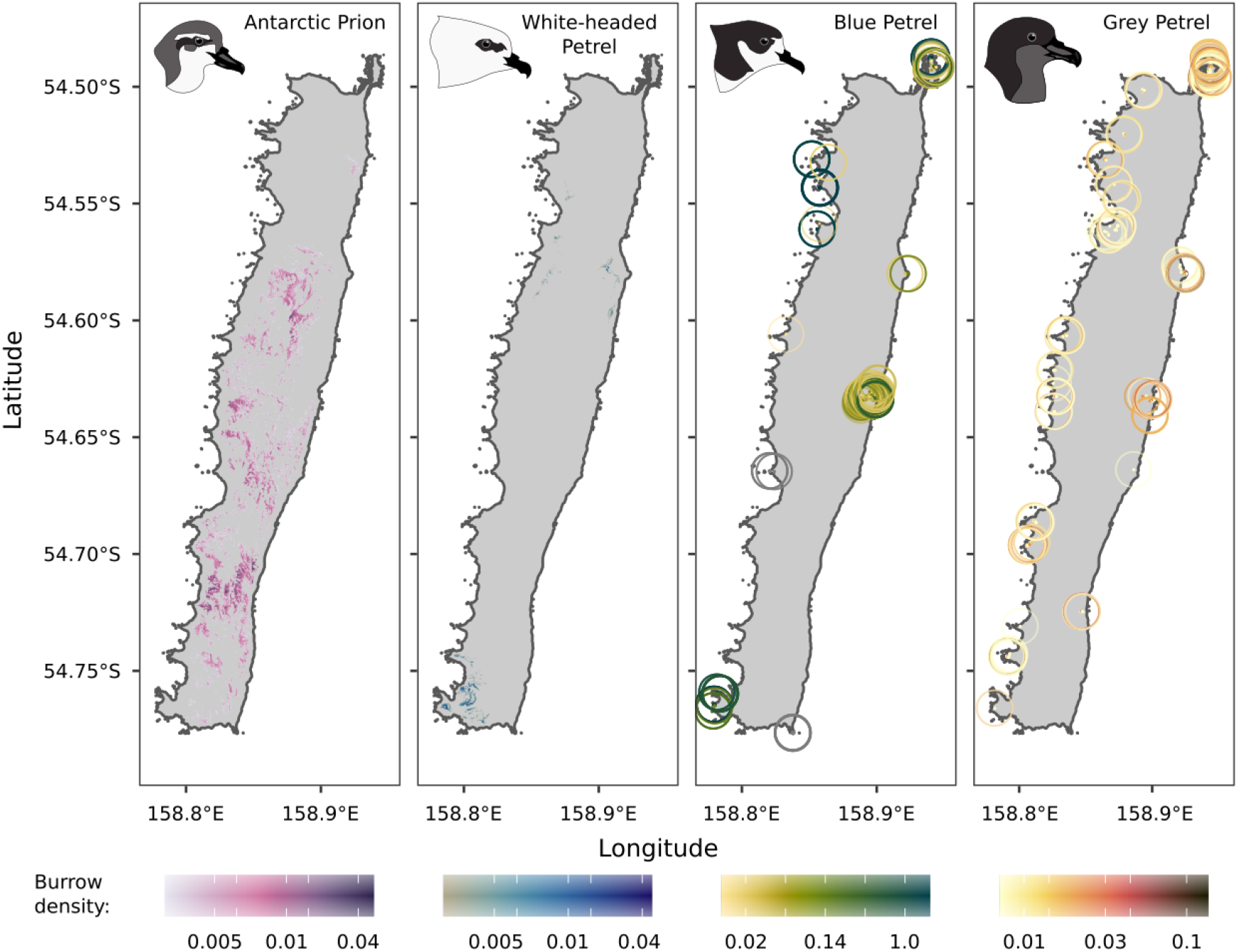
Island-wide burrow density plotted on a log scale for widespread species a) Antarctic Prions and b) White-headed Petrels, and recolonising species c) Blue Petrels and d) Grey Petrels. a) and b) are DSM predictions, c) and d) are from search surveys—circles are plotted around occupied pixels to increase visibility. Predicted densities in a) and b) were truncated below 1% of peak predicted density.

### Comparing methods

Aside from two Blue Petrel burrows, only Antarctic Prions and White-headed Petrels were recorded during stratified random island-wide surveys. Prions were found in 13% of plots and White-headed Petrels in <3%. Transects yielded dramatically higher sample sizes. Antarctic Prions were an order of magnitude more common than other species and as a result, distance analysis of prion data from transects resulted in confidence intervals just over one quarter of the width of those from plots, and one third the width of those for White-headed Petrels (Table 1). Uncertainty was higher overall for the less abundant White-headed Petrels, with negligible difference between the design- and model-based methods for estimating burrow numbers. DSMs resulted in marginally higher precision in estimates for both prions and White-headed Petrels, based either on the delta or variance propagation method for determining uncertainty.

Targeted surveys resulted in similar apparent precision (based upon CV) of estimated Blue Petrel and Grey Petrel burrow numbers, as the randomised surveys achieved for Antarctic Prions (Table 1). Model-based estimates appear to substantially over-predict current Blue and Grey Petrel burrow-densities, resulting in estimates two to three orders of magnitude above our direct estimates (Figure 3). We overlaid our searched area—buffered nocturnal, transect and search survey tracks—on modelled Blue and Grey Petrel abundance. This comparison indicates we covered habitat supporting 30% and 18% of the model-predicted populations of each species, confirming the species were indeed absent from those areas, and that the models are over-predicting (Figure 3).

**Figure 3:**
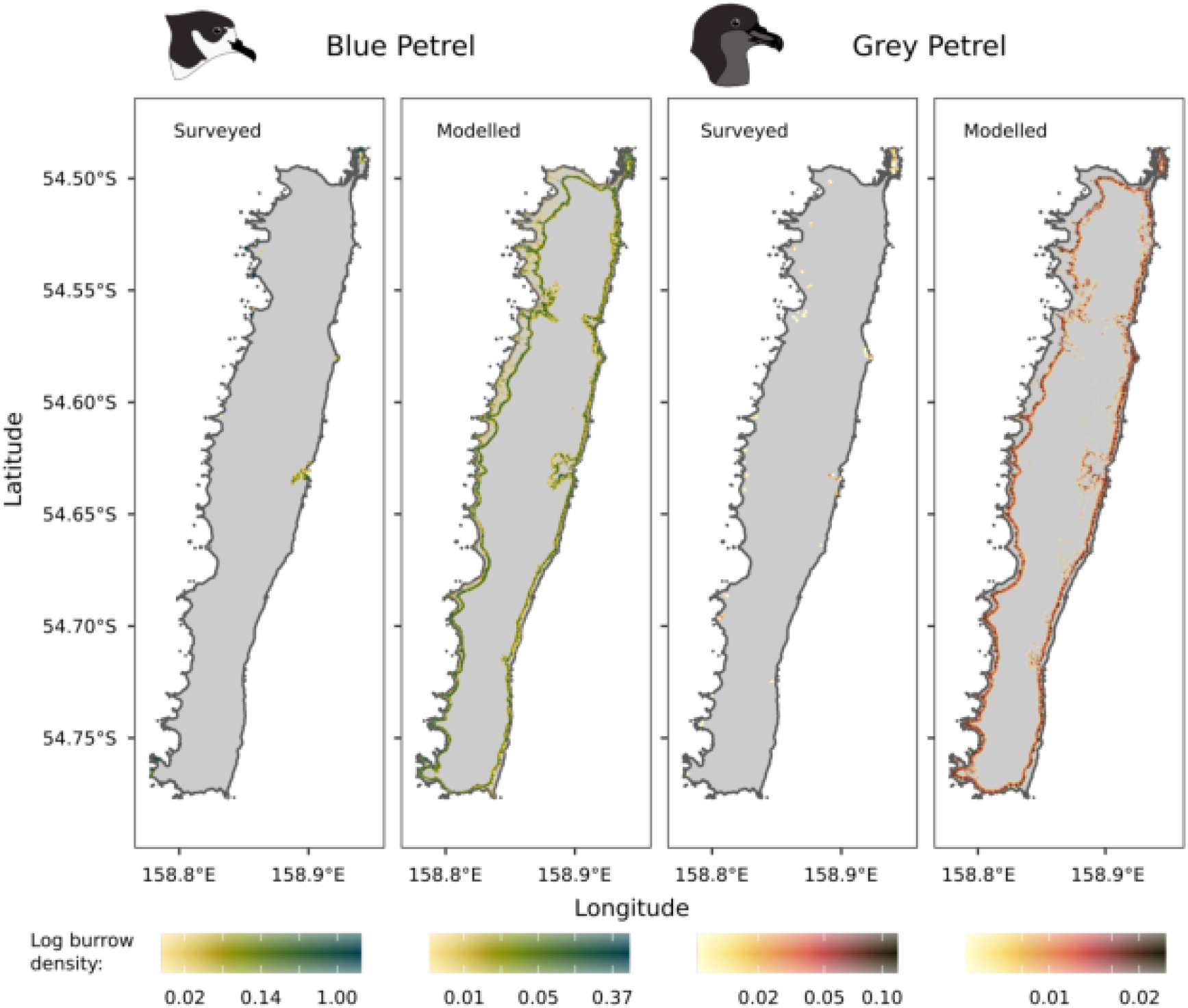
Surveyed and modelled densities of recolonising Blue Petrels and Grey Petrels illustrating that models currently over-predict the distributions of both species.

## DISCUSSION

Our results clearly show that a multi-method approach is needed to study petrel responses following a large-island invasive species eradication, owing to the divergent ecologies and life histories of the species. Surveys cannot be based upon only what managers think they know, or assumed knowledge, before eradication. They must be flexible enough to detect novel changes, and simultaneously deal with very common and very rare species. By combining unbiased and targeted survey methods, and design-based and model-based analyses we were able to collect information to detect quantitative changes in abundance occurring in widespread species, and qualitative changes such as colonisation events in rare species, and at new locations that are not easily predictable.

Importantly, our results show that the two recolonising species remain highly localised, while the two more widespread species, White-headed Petrels and Antarctic Prions, continue to occur at abundances well short of apparent historic maxima based upon early descriptions (Cumpston 1968). Our results show the distribution of White-headed Petrels appears to have contracted since the 1970s (Warham 1967, Cumpston 1968, Brothers 1984). Search surveys located Blue Petrels in 0.05% and Grey Petrels in 0.03% of 20 × 20m pixels island-wide, while models predict >1 breeding pair of Antarctic Prions in 16% and White-headed Petrels in 0.9% pixels. Although perhaps not surprising given the short time since eradication, these results highlight that the island is in the earliest stages of recovery and provide a useful benchmark for measuring future change in species distributions.

Over 80% of petrel population estimates are collected with a view to informing status and trend assessments (Bird et al. submitted A). With such objectives, unbiased survey designs are important (Buckland et al. 2015). However, our results highlighted the important influence of species rarity on these models, which resulted in relatively poor performance, and failed to restrict confidence intervals to reasonable limits for detecting trends for all but the most widespread species, Antarctic Prion. Distance-sampling along transects was a far more time-efficient and precise way of collecting spatially explicit abundance data.

In theory our staged approach to survey design lends itself to combining model-based analyses and survey design by allowing for extra effort to be included in areas with known detections while permitting statistically rigorous estimates (Pacifici et al. 2016). The utility of this approach in reality is unclear, given we recorded so few burrows in our whole-island transects that were not White-headed Petrels or Antarctic Prions. Future surveys of this nature could look at improving and developing a more balanced stratified survey design by generating presence-only model-based predictions of occurrence using preliminary survey data, including known sites. However, had we sought a more balanced design in our case there is a risk we would still have failed to record rare species while also collecting less data on widespread species. The success of our transect survey for informing estimates of White-headed Petrel and particularly Antarctic Prion populations and distributions can be attributed to the preliminary survey which allowed us to weight survey effort by stratum. In this context, data-driven survey design proved effective.

While targeted searches resulted in high precision in burrow estimates, this may be at the expense of accuracy. An untested assumption in most search surveys is that all, or most colonies are found. Without a randomised survey design, it is not possible to assess what may have been missed. Uncertainty in the assumptions of search-based surveys means it would be difficult to use repeat surveys alone to infer numeric changes in the population, but the distributional information collected would provide a valuable baseline for future inference.

We found spotlighting to be a highly effective part of our survey strategy. Having only encountered three species’ burrows during daytime transect surveys, we encountered flying/vocalising birds of 11 species while spotlighting at night, which led to the subsequent discovery of nesting burrows of three of these additional species during follow-up daytime searches.

Given the high uncertainty in most design-based petrel population assessments to date (Bird et al. submitted A), one of our aims was to assess whether model-based approaches could improve precision in population estimates. While uncertainty in estimated prion burrow numbers was at the lower end of published estimates for petrels, combining variance in occupancy estimates when generating adjusted population estimates still resulted in a level of overall uncertainty that is likely to limit the detection of significant trends from repeated surveys (Bird et al. submitted A). We found that model-based methods yielded marginally narrower confidence intervals for prions than typical distance analysis, a hybrid of design- and model-based methods (Camp et al. 2020). Importantly, there appears to be a threshold in terms of how abundant or uniformly distributed a species is which will govern first whether an unbiased or targeted survey design should be used, and second whether design-based or model-based analysis is preferable. For example, White-headed Petrels were rarer than Antarctic Prions and our methods delivered lower precision overall, with negligible difference between them. The fact that model-based methods consistently estimated lower burrow numbers than design-based methods for Antarctic Prions and White-headed Petrels suggests they were not over-predicting, and therefore the models were a reasonable approximation of the realized niches of both species (Rayner et al. 2007, Camp et al. 2020).

To test the assumption that our search surveys had successfully detected most Blue and Grey Petrel colonies we overlaid our searched area, based upon buffered search tracks, on top of modelled abundance. Despite targeting field searches on potentially suitable habitat our search area covered only 30% of total model-predicted Blue Petrel burrows and 18% of predicted Grey Petrel burrows. One possibility is that our searches missed a substantial proportion of these recolonising populations. However, field rangers have been employed continuously on Macquarie Island since the late 1990s when both species began to recolonise. There has been a considerable focus by the reserve manager (Tasmanian Parks and Wildlife Service) to record the locations of burrowing petrels, including designated burrowing petrel tasks over much of that time, resulting in gradual discovery of new petrel breeding sites. In the context of this effort, together with our dedicated searches, it seems unlikely that a significant part of either population was missed. More likely, our models failed to capture these species’ realized niches accurately. For highly localised species, the models were unable to accurately discriminate between suitable occupied habitat and apparently suitable unoccupied habitat. We were able to generate a large number of real absences from our combined datasets to use in models. However, initial attempts to run models with 30,000 absences failed computationally, so a sub-sampled set of 3,000 absences was used in the final models. Given their rarity in the landscape, much apparently suitable habitat must still exist outside of real and modelled absences. It is also possible that our models were too simplistic and failed to capture some important predictor of occurrence such as proximity to a headland.

Beyond population estimation, density surface models of species distribution and habitat suitability have wider applications. For example, the apparent over-prediction of Blue and Grey Petrels by our GAMs suggests we could employ presence-only models to predict suitable habitat island-wide (Cianfrani et al. 2010). Although future population increase will be contingent on external factors such as regional population trends, proximity to source populations, and marine resources (Ashmole 1963, Brooke et al. 2018), suitable habitat models could provide an indication of the potential for long-term growth. By providing information on distribution as well as abundance, models can be more appropriate for communicating information to non-experts than the results of unbiased survey designs alone (Miller et al. 2013). Species distribution maps provide a useful benchmark for assessing future change and can inform monitoring designed to detect future population expansion.

Constant monitoring within established long-term plots is a reliable way of detecting population change, but when densities reach carrying capacity within monitored colonies, populations will expand into unoccupied habitat (Kildaw et al. 2005). To detect such expansion model-predicted suitable habitat can be used to select additional plots in currently unoccupied areas.

### Monitoring implications

To date, whole-island surveys of petrel populations have not been successful in generating time-stamped data from which trends can be detected, largely owing to their inherent uncertainties (Bird et al. submitted A). Our study provides further evidence that whole-island surveys, regardless of method, yield wide confidence limits when used to provide population estimates, which in turn may preclude their use for reliably detecting trends. Instead, if the primary goal is to measure population change, approaches that focus on monitoring more frequently within fixed plots are likely to be more accurate (Buxton et al. 2016).

The disproportionate number of petrels that are threatened, together with a societal focus on threatened species as conservation priorities, means that detecting trends to inform conservation assessments commonly is the main goal for petrel monitoring. Rarity of threatened species adds to the already considerable challenges of surveying burrowing seabirds, and there is an argument that the additional costs associated with rare species surveys in terms of high uncertainty and untestable assumptions outweigh the potential benefits of the data. However, there are other reasons to monitor seabirds and measure and map populations. We highlight the utility of species population estimates and their distributions for communicating outcomes, guiding on-island management, and benchmarking distributions to assess ecological function and potential future expansion, particularly when spatially explicit predictions of abundance are provided through model-based analyses.

Whatever the end-use of a population estimate it is preferable to maximise accuracy and precision. Here we’ve shown that a novel application of a density surface model for a common widespread burrowing seabird yielded a relatively precise population estimate. However, unbiased survey designs and model-based analyses, which are typically the most accurate, performed poorly for rare, localised species. In these cases, targeted surveys are needed, despite their inherent assumptions. To maximise accuracy and precision across a diverse assemblage that includes both common widespread species and rare localised species, a diversity of approaches is required. Rather than focussing efforts on threatened species recovery looking at common and widespread species provides information on the restoration of ecological function as supported by abundant seabird populations. Unlike with rare species, we found that unbiased survey designs and model-based analyses for a common widespread species were more robust to design and model assumptions. Given favourable recovery trajectories even currently rare species will hopefully be more readily surveyable using unbiased survey designs in the future, particularly with a greater focus on combining model-predictions in survey design (Pacifici et al. 2016).

Importantly, the data collected during this study will improve understanding of ecological responses to invasive species eradication. Globally, reporting of seabird responses to eradications has been rare (Brooke et al. 2018, Towns 2018), and no other examples of multi-species responses to predator removal exist at the scale of Macquarie Island. With several decades of data now available, Macquarie Island provides a unique case study.

## Notes

### Competing Interest Statement

The authors have declared no competing interest.

