## Supporting Information for "Detecting seabird responses to invasive species eradication"

### Appendix S1: Supporting methods.

***ii) Environmental covariates, stratification and survey design***

We stratified the island and used the distribution information collected during our preliminary survey to assign survey effort preferentially to strata with higher burrow encounter rates to minimise uncertainty around higher density estimates (Arneill et al. 2019). We used seven spatial environmental covariates in an unsupervised fuzzy c-means cluster analysis (Puspini 1969, Bezdek 1980) to define eight strata across the island. The environmental covariates were aspect, elevation, ridgeness, slope, a topographic wetness index, topographically-deflected mean wind speed, and a normalised difference vegetation index (NDVI). They were derived from a 5 m resolution digital elevation model and QuickBird satellite imagery (Bricher et al. 2013). All burrow locations were overlain on the eight strata and classified by species and stratum (Figure 1). The observed frequency of each species’ burrows by stratum was compared with expected frequency derived from track points extracted from our GPS tracks in Garmin BaseCamp version 4.7.0 (Garmin 2019)—a proxy for time spent surveying each stratum.

Randomised survey designs perform well for widespread, dispersed species but poorly for highly localised, clustered populations (Dilley et al. 2019). Therefore, our survey focussed on White-headed Petrels and Antarctic Prions, the two most widespread and abundant species during our preliminary survey, and previous surveys (Brothers 1984, Brothers and Bone 2008). The observed and expected frequency distributions (Figure 1) were used to generate 200 stratified random points and 100 reserve points (as replacements when a target point was inaccessible) in QGIS v3.6.3 (QGIS Development Team 2019). One third (67 × 2) of the points were assigned to strata based upon the observed frequencies of Antarctic Prion and White-headed Petrel burrows in different strata, and the remaining third (66) were controls based upon the relative extents of the eight strata island-wide.

***iii) Survey implementation***

*Island-wide plot and transect surveys*

A simulation study found that in most cases >100 plots are required to attain a power of 0.8 to detect changes between numbers of breeding petrels between repeat surveys, and that power increases considerably when plot size increases from 3 m to 5 m radius (Buxton et al. 2016). From 5^th^ January to 24^th^ April 2018 we measured burrow density at 194 of the randomly generated points. At each sample location we thoroughly searched 100 m^2^ within a circle of radius 5.66 m (measured with a length of rope). All burrows were identified to species (see Bird et al. submitted B), counted and recorded by handheld GPS if the centre of the burrow entrance was within the plot.

To generate a second measure of burrow density we recorded burrows along 158 km of transects navigated between plots. When complete searches of the sampling area are not possible survey design must overcome imperfect detection of burrows (Mackenzie and Royle 2005). Distance sampling is a way of estimating density while explicitly accounting for uncertain detection (Miller et al. 2013), and has been used effectively to estimate petrel population sizes (Lawton et al. 2006, Rexer-Huber et al. 2017). In our survey, a lead person walked the shortest navigable distance in an approximately straight line, recording the exact route by handheld GPS. Waypoints were taken each time the transect transitioned from ‘short’ to ‘tall’ (</>70 cm) vegetation so that the transect could be subdivided post-hoc for analyses. A second person, walking directly behind the first, visually scanned left and right identifying all burrows observed from the transect line, recording them on a second GPS, and measuring the perpendicular distance between the burrow and the transect line to the nearest 10 cm.

*Targeted species-specific searches*

For species not encountered incidentally during the preliminary survey, i.e. they were not widespread away from known colonies, we undertook targeted searches. In 2017 and 2018 we surveyed all known colonies of Blue Petrels and Grey Petrels. To detect new colonies we undertook targeted searches following methods adapted from Dilley *et al*. (2017) and Schulz *et al*. (2006). We defined suitable habitats based upon descriptions in these two papers and our own experience of known sites (see Bird et al. submitted B). We conducted nocturnal surveys of suitable habitat by observing from vantage points, or traversing on foot, to identify birds vocalising or in flight attending colonies. Flying birds were readily observed at >100 m range with a Ledlenser P17.2 spotlight. Active daytime burrow searches were made in suitable habitats focusing on areas where Blue Petrel or Grey Petrel burrows were recorded during island-wide transects, any areas where birds were recorded during nocturnal surveys, and areas around Brown Skua *Stercorarius antarcticus* territories where Blue Petrel remains were identified.

Burrows were found by investigating dark-green patches of tussock grass (*Poa foliosa* and/or *Poa cookii*) or *Acaena* *magellanica* (caused by the combined manuring effect of a concentration of birds), by looking for fresh droppings, feathers and trampled vegetation or soil at burrow entrances, and by listening for Blue Petrels responding to imitation calls (Schulz et al. 2006, Dilley et al. 2017). Colonies were defined as clusters of ≥1 burrow separated by >50 m from the nearest cluster.

We attempted to census Grey Petrels, searching outwards from all located burrows to search the whole area of suitable habitat (Schulz et al. 2006). When surveying Blue Petrel colonies, we followed Dilley et al. (2017).

Arneill, G. E. et al. 2019. Sampling strategies for species with high breeding-site fidelity: A case study in burrow-nesting seabirds (C Lebarbenchon, Ed.). - PLOS ONE 14: e0221625.

Bezdek, J. C. 1980. A Convergence Theorem for the Fuzzy ISODATA Clustering Algorithms. - IEEE Trans. Pattern Anal. Mach. Intell. PAMI-2: 1–8.

Bricher, P. K. et al. 2013. Mapping sub-Antarctic cushion plants using random forests to combine very high resolution satellite imagery and terrain modelling. - Plos One 8: e72093.

Brothers, N. P. 1984. Breeding, Distribution and Status of Burrow-nesting Petrels at Macqaurie Island. - Aust. Wildl. Res. 11: 113–131.

Brothers, N. and Bone, C. 2008. The response of burrow-nesting petrels and other vulnerable bird species to vertebrate pest management and climate change on sub-Antarctic Macquarie Island. - Pap. Proc. R. Soc. Tasman. 142: 123–148.

Buxton, R. T. et al. 2016. Monitoring burrowing petrel populations: A sampling scheme for the management of an island keystone species. - J. Wildl. Manag. 80: 149–161.

Dilley, B. J. et al. 2017. The distribution and abundance of Blue Petrels (Halobaena caerulea) breeding at subantarctic Marion Island. - Emu-Austral Ornithol.: 1–11.

Dilley, B. J. et al. 2019. Clustered or dispersed: testing the effect of sampling strategy to census burrow-nesting petrels with varied distributions at sub-Antarctic Marion Island. - Antarct. Sci.: 1–12.

Garmin 2019. BaseCamp.

Lawton, K. et al. 2006. An estimate of population sizes of burrowing seabirds at the Diego Ramirez archipelago, Chile, using distance sampling and burrow-scoping. - Polar Biol. 29: 229–238.

Mackenzie, D. I. and Royle, J. A. 2005. Designing occupancy studies: general advice and allocating survey effort. - J. Appl. Ecol. 42: 1105–1114.

Miller, D. L. et al. 2013. Spatial models for distance sampling data: recent developments and future directions. - Methods Ecol. Evol. 4: 1001–1010.

Puspini, E. H. 1969. A New Approach to Clustering. - Inf. Control 19: 22–32.

QGIS Development Team 2019. QGIS Geographic Information System.

Rexer-Huber, K. et al. 2017. White-chinned petrel population estimate, Disappointment Island (Auckland Islands). - Polar Biol. 40: 1053–1061.

Schulz, M. et al. 2006. Breeding of the Grey Petrel (Procellaria cinerea) on Macquarie Island: population size and nesting habitat. - Emu 105: 323–329.

Appendix S2: Distance sampling – detection function selection and diagnostics

**Table S2-1:** Detection function selection for Antarctic Prions. Following Howe et al. (2019) we used an omnibus overdispersion factor (ĉ). The same value of ĉ was used for all models with the same key function. It was derived from a χ2 GOF test of the most highly parameterised in each key function divided by its degrees of freedom (df). ĉ was used in the calculation of adjusted AIC (QAIC). QAIC was used to select the preferred model from each key function (half-normal and hazard-rate). Of the two candidates the model with the lowest value when the χ2 GOF test statistic was divided by df was selected – a hazard-rate model with no adjustments. Candidate models are highlighted in bold, the selected model all in bold.

| df | AIC | χ2 GOF/df | P.chisq | key | id | ĉ | QAIC |
| --- | --- | --- | --- | --- | --- | --- | --- |
| 1 | 7585.746 | **11.28206** | 0 | ds_hn | ds_hn0 | 11.42716 | **665.6599** |
| 2 | 7578.059 | 10.95111 | 0 | ds_hn | ds_hncos2 | 11.42716 | 666.8123 |
| 3 | 7569.836 | 10.70283 | 0 | ds_hn | ds_hncos3 | 11.42716 | 667.9176 |
| 4 | 7571.708 | 10.91875 | 0 | ds_hn | ds_hncos4 | 11.42716 | 669.9064 |
| 6 | 25665.71 | 5.72E+08 | 0 | ds_hn | ds_hncos6 | 11.42716 | 2256.978 |
| 2 | 7584.734 | 11.38588 | 0 | ds_hn | ds_hnherm2 | 11.42716 | 667.3964 |
| 3 | 7563.796 | 10.47612 | 0 | ds_hn | ds_hnpoly2 | 11.42716 | 667.389 |
| 5 | 7567.587 | 10.92712 | 0 | ds_hn | ds_hnpoly4 | 11.42716 | 671.3708 |
| 7 | 7571.535 | 11.42716 | 0 | ds_hn | ds_hnpoly6 | 11.42716 | 675.3663 |
| **2** | **7560.039** | **10.22083** | **0** | **ds_hr** | **ds_hr0** | **11.609** | **654.8775** |
| 3 | 7562.049 | 10.44286 | 0 | ds_hr | ds_hrcos2 | 11.609 | 656.8783 |
| 4 | 7563.975 | 10.69473 | 0 | ds_hr | ds_hrcos3 | 11.609 | 658.872 |
| 5 | 7565.227 | 10.85643 | 0 | ds_hr | ds_hrcos4 | 11.609 | 660.8075 |
| 3 | 7562.049 | 10.44286 | 0 | ds_hr | ds_hrherm2 | 11.609 | 656.8783 |
| 4 | 7563.384 | 10.61812 | 0 | ds_hr | ds_hrpoly2 | 11.609 | 658.8211 |
| 6 | 7565.625 | 11.08561 | 0 | ds_hr | ds_hrpoly4 | 11.609 | 662.6696 |
| 8 | 7569.571 | 11.609 | 0 | ds_hr | ds_hrpoly6 | 11.609 | 666.6648 |

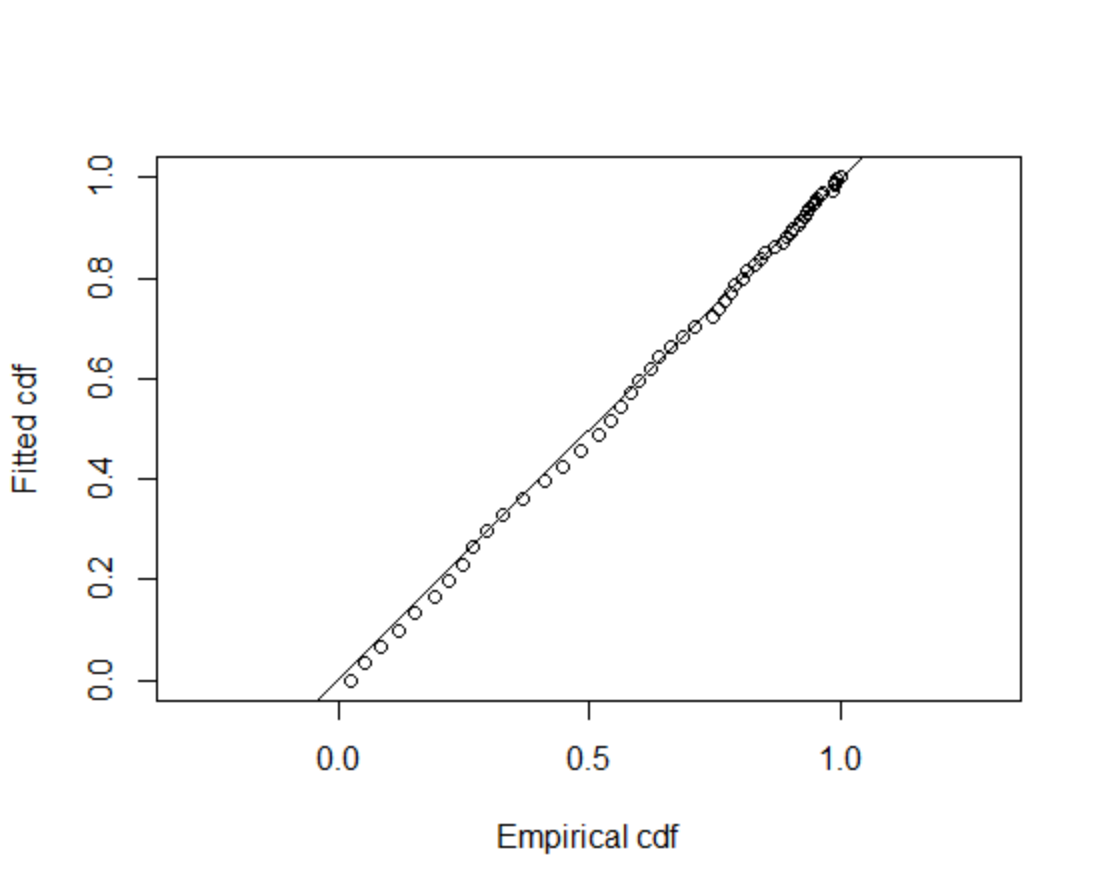

**Figure S2-1:** Quantile-quantile plot for the detection function model selected to estimate Antarctic Prion burrow detection probability. The ﬁtted cumulative distribution function (cdf) is plotted against the empirical cdf. The points seem to fall about the straight line, which provides evidence the function adequately ﬁts the data.

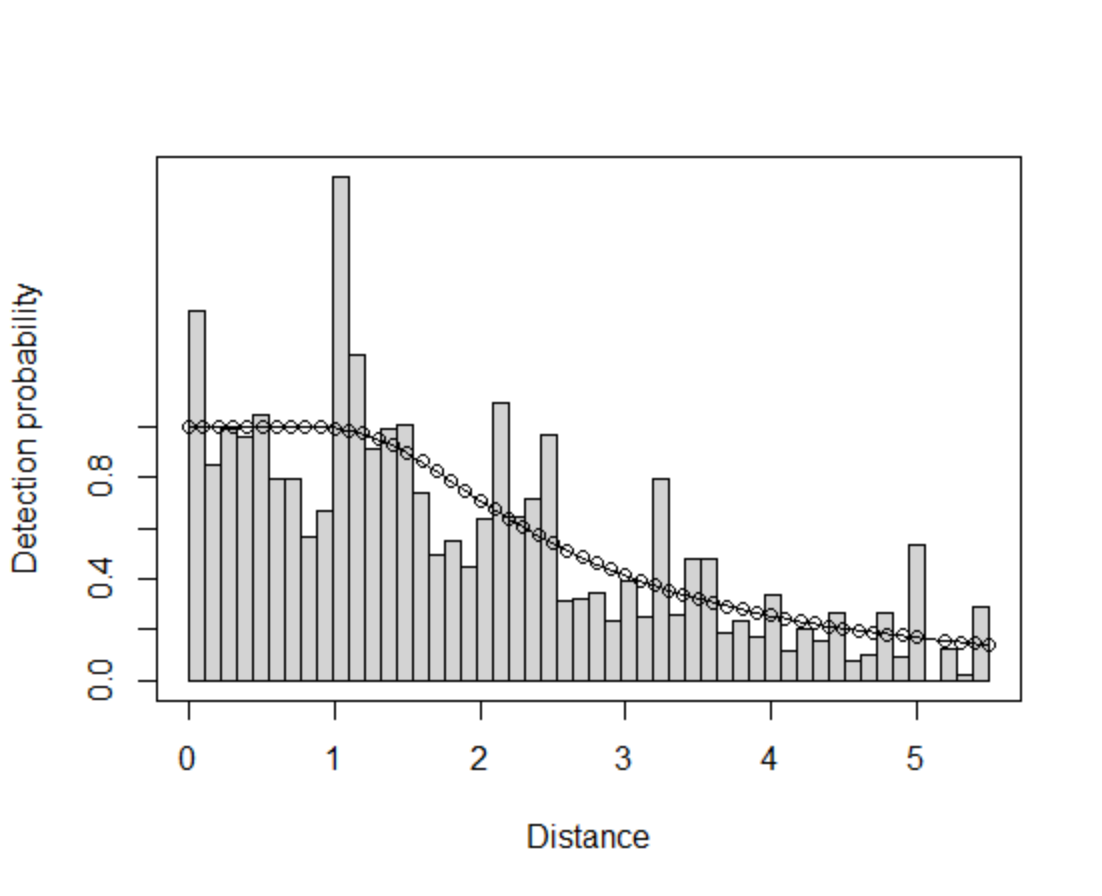

**Figure S2-2:** Detection probability for the model selected for Antarctic Prion burrows.

**Table S2-2:** Detection function selection for White-headed Petrels. Following Howe et al. (2019) we used an omnibus overdispersion factor (ĉ). The same value of ĉ was used for all models with the same key function. It was derived from a χ2 GOF test of the most highly parameterised in each key function divided by its degrees of freedom (df). ĉ was used in the calculation of adjusted AIC (QAIC). QAIC was used to select the preferred model from each key function (half-normal and hazard-rate). Of the two candidates the model with the lowest value when the χ2 GOF test statistic was divided by df was selected – a half-normal model with observer as a covariate. Candidate models are highlighted in bold, the selected model all in bold.

| df | AIC | χ2 GOF/df | P.chisq | key | id | ĉ | QAIC |
| --- | --- | --- | --- | --- | --- | --- | --- |
| 1 | 1064.137 | 3.274495 | 1.68E-05 | ds_hn | ds_hn0 | 2.584583 | 412.951 |
| 2 | 1057.63 | 1.925359 | 0.019514 | ds_hn | ds_hncos2 | 2.584583 | 411.6596 |
| 3 | 1055.261 | 1.839396 | 0.031948 | ds_hn | ds_hncos3 | 2.584583 | 411.9692 |
| 4 | 1055.804 | 1.849116 | 0.035451 | ds_hn | ds_hncos4 | 2.584583 | 413.4052 |
| 6 | 1058.684 | 2.122043 | 0.019608 | ds_hn | ds_hncos6 | 2.584583 | 416.972 |
| 2 | 1065.773 | 3.431417 | 1.29E-05 | ds_hn | ds_hnherm2 | 2.584583 | 414.81 |
| **2** | **1015.871** | **2.027202** | **0.012659** | **ds_hn** | **ds_hnobs** | **2.584583** | **395.5027** |
| 3 | 1054.246 | 1.807733 | 0.036048 | ds_hn | ds_hnpoly2 | 2.584583 | 411.5762 |
| 5 | 1058.154 | 2.127002 | 0.015529 | ds_hn | ds_hnpoly4 | 2.584583 | 415.5408 |
| 7 | 1062.085 | 2.584583 | 0.005635 | ds_hn | ds_hnpoly6 | 2.584583 | 419.514 |
| 2 | 1058.036 | 2.118966 | 0.008483 | ds_hr | ds_hr0 | 3.287334 | 324.6354 |
| 3 | 1054.425 | 1.791247 | 0.038367 | ds_hr | ds_hrcos2 | 3.287334 | 324.9286 |
| 4 | 1056.422 | 1.938769 | 0.025557 | ds_hr | ds_hrcos3 | 3.287334 | 326.9278 |
| 5 | 1056.915 | 2.013762 | 0.023226 | ds_hr | ds_hrcos4 | 3.287334 | 328.4692 |
| 7 | 1059.725 | 2.316778 | 0.013329 | ds_hr | ds_hrcos6 | 3.287334 | 332.1074 |
| 3 | 1059.759 | 2.250531 | 0.006018 | ds_hr | ds_hrherm2 | 3.287334 | 326.5513 |
| 3 | 1015.555 | **2.074022** | 0.01259 | ds_hr | ds_hrobs | 3.287334 | **313.1045** |
| 4 | 1063.78 | 3.22629 | 0.000117 | ds_hr | ds_hrpoly2 | 3.287334 | 329.1659 |
| 6 | 1063.816 | 2.807012 | 0.001759 | ds_hr | ds_hrpoly4 | 3.287334 | 331.9601 |
| 8 | 1066.234 | 3.287334 | 0.000934 | ds_hr | ds_hrpoly6 | 3.287334 | 335.4791 |

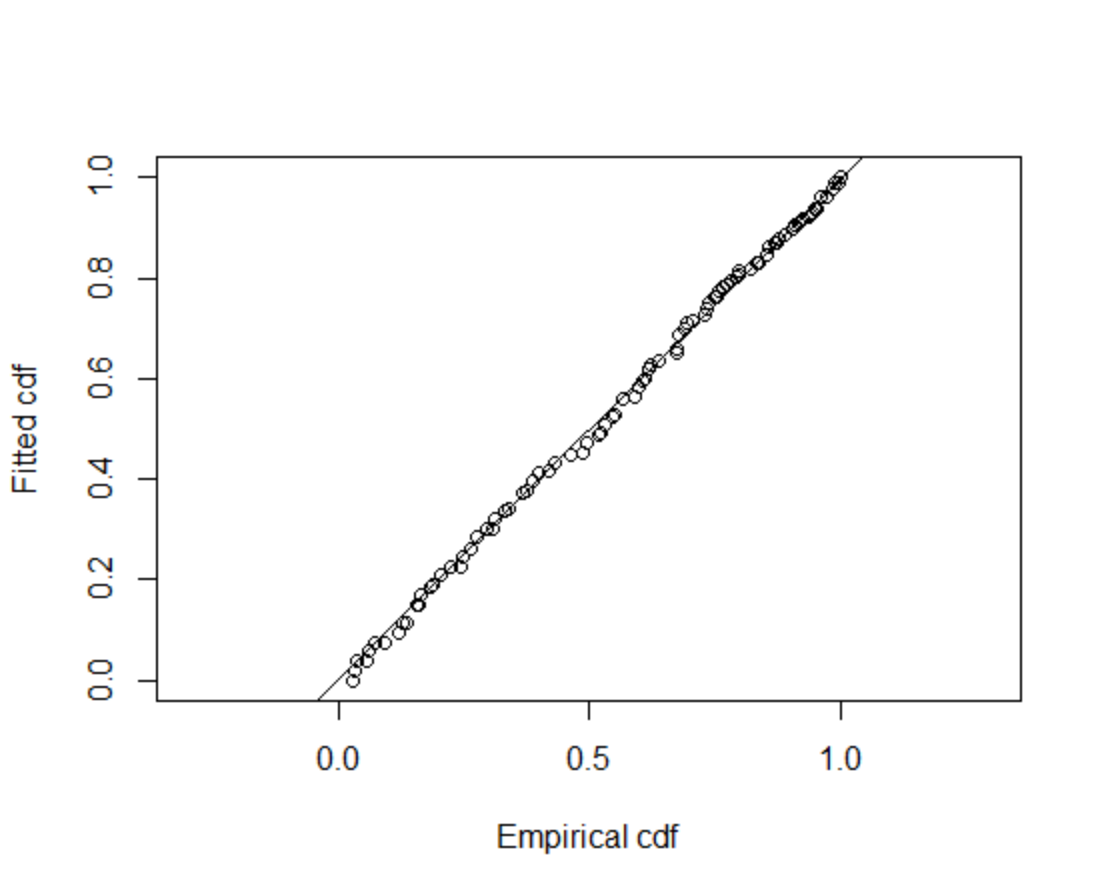

**Figure S2-3:** Quantile-quantile plot for the detection function model selected to estimate White-headed Petrel burrow detection probability. The ﬁtted cumulative distribution function (cdf) is plotted against the empirical cdf. The points seem to fall about the straight line, which provides evidence the function adequately ﬁts the data.

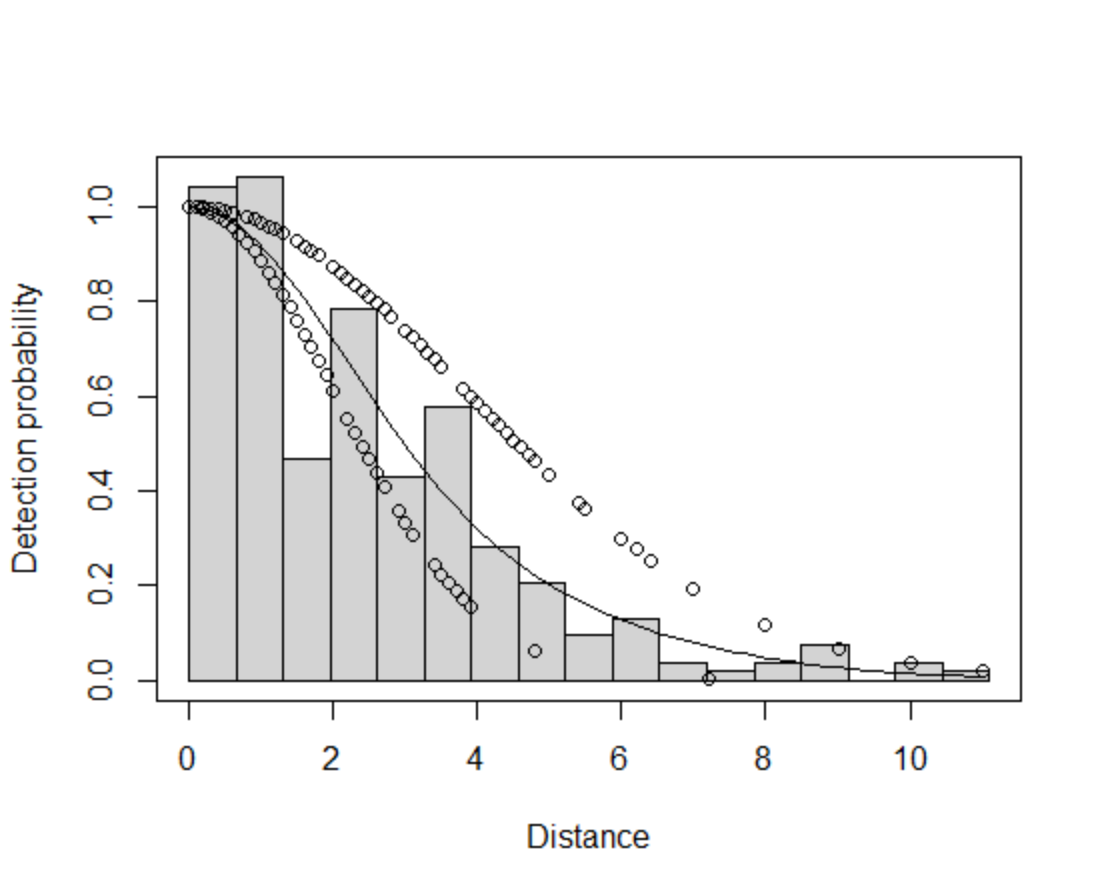

**Figure S2-4:** Detection probability for the model selected for White-headed Petrel burrows.

Appendix S3: Blue Petrel colony estimates

**Table S3-1:** Population density derived from 1 and 2 m radius plots sampled within Blue Petrel colonies. See Dilley et al. (2017) for full methods.

| **site** | **Veg type** | **Density (brws/m^2^)** | **n plots** | **sd** | **area m^2^** | **est** |
| --- | --- | --- | --- | --- | --- | --- |
| Caroline Point A | Tussock | 0.59 | 13 | 0.56 | 1036 | 609 |
| Caroline Point B | Colobanthus | 0.99 | 1 | 0.00 | 136 | 135 |
| Douglas Point stack | Tussock | 1.61 | 24 | 0.65 | 1321 | 2121 |
| Green Gorge GP-188 | Tussock | 0.41 | 24 | 0.45 | 1022 | 421 |
| North Head GP-54 | Tussock | 1.73 | 21 | 0.69 | 724 | 1251 |
| North Head GP-55 | Tussock | 1.79 | 18 | 0.41 | 720 | 1287 |
| North Head GP-56 | Tussock | 1.41 | 19 | 0.82 | 607 | 855 |
| North Head GP-57 | Tussock | 0.88 | 9 | 0.88 | 671 | 593 |
| Langdon Point stack | Tussock | 1.29 | 16 | 0.75 | 480 | 621 |
| Mawson Point main stack | Tussock | 1.49 | 6 | 0.96 | 247 | 367 |
| Tottan Head B | Tussock | 0.28 | 11 | 0.28 | 543 | 153 |
| Upper Goat Bay | Tussock | 0.24 | 4 | 0.09 | 167 | 40 |
| West Rock north | Colobanthus | 1.19 | 26 | 0.84 | 1547 | 1838 |

Appendix S4: Density surface model selection

**Table S4-1:** Model selection for Antarctic Prions. Diagnostic plots for global models were compared for three distributions. Then the highest non-significant covariate was dropped until there was no improvement in AIC.

| **model** | **response** | **terms** | **adjusted *R^2^*** | **AIC** | **REML** | **deviance explained** |
| --- | --- | --- | --- | --- | --- | --- |
| dsm_ap_nb3 | negative binomial | s(x,y), s(dem), s(ndvi), s(slope) | 0.11 | 8005.25 | 4038.67 | 46.28% |
| dsm_ap_nb1 | negative binomial | s(x,y), s(dem), s(ndvi), s(ridge), s(slope), s(wetness) | 0.11 | 8011.05 | 4041.37 | 46.38% |
| dsm_ap_nb | negative binomial | s(x,y), s(dem), s(ndvi), s(ridge), s(slope), s(wetness), s(wind) | 0.11 | 8012.09 | 4043.26 | 46.41% |
| dsm_ap_nb2 | negative binomial | s(x,y), s(dem), s(ndvi), s(slope) s(wetness) | 0.11 | 8013.09 | 4040.12 | 46.31% |
| dsm_ap_tw | tweedie | s(x,y), s(dem), s(ndvi), s(ridge), s(slope), s(wetness), s(wind) | 0.13 | 16576.84 | 4039.38 | 40.89% |
| dsm_ap_qp | quasipoisson | s(x,y), s(dem), s(ndvi), s(ridge), s(slope), s(wetness), s(wind) | 0.16 | NA | 4238.82 | 39.19% |

```
Family: Negative Binomial(0.103)
Link function: log

Formula:
count ~ s(x, y, k = 100) + s(dem, k = 10) + s(ndvi, k = 10) +
    s(slope, k = 10) + offset(off.set)

Parametric coefficients:
            Estimate Std. Error z value Pr(>|z|)
(Intercept)   -9.104      0.317  -28.72   <2e-16 ***
---
Signif. codes:  0 ‘***’ 0.001 ‘**’ 0.01 ‘*’ 0.05 ‘.’ 0.1 ‘ ’ 1

Approximate significance of smooth terms:
            edf Ref.df Chi.sq  p-value
s(x,y)   46.851 59.758 282.12  < 2e-16 ***
s(dem)    6.203  7.262  93.24 2.29e-16 ***
s(ndvi)   2.066  2.626 365.35  < 2e-16 ***
s(slope)  3.732  4.707  22.50 0.000323 ***
---
Signif. codes:  0 ‘***’ 0.001 ‘**’ 0.01 ‘*’ 0.05 ‘.’ 0.1 ‘ ’ 1

R-sq.(adj) =  0.107   Deviance explained = 46.3%
-REML = 4038.7  Scale est. = 1         n = 14118

Method: REML   Optimizer: outer newton
full convergence after 7 iterations.
Gradient range [-6.374027e-06,5.004075e-06]
(score 4038.669 & scale 1).
Hessian positive definite, eigenvalue range [0.2206379,311.5018].
Model rank =  127 / 127

Basis dimension (k) checking results. Low p-value (k-index<1) may
indicate that k is too low, especially if edf is close to k'.

            k'   edf k-index p-value
s(x,y)   99.00 46.85    0.59  <2e-16 ***
s(dem)    9.00  6.20    0.67  <2e-16 ***
s(ndvi)   9.00  2.07    0.74   0.015 *
s(slope)  9.00  3.73    0.76   0.440
---
Signif. codes:  0 ‘***’ 0.001 ‘**’ 0.01 ‘*’ 0.05 ‘.’ 0.1 ‘ ’ 1
```

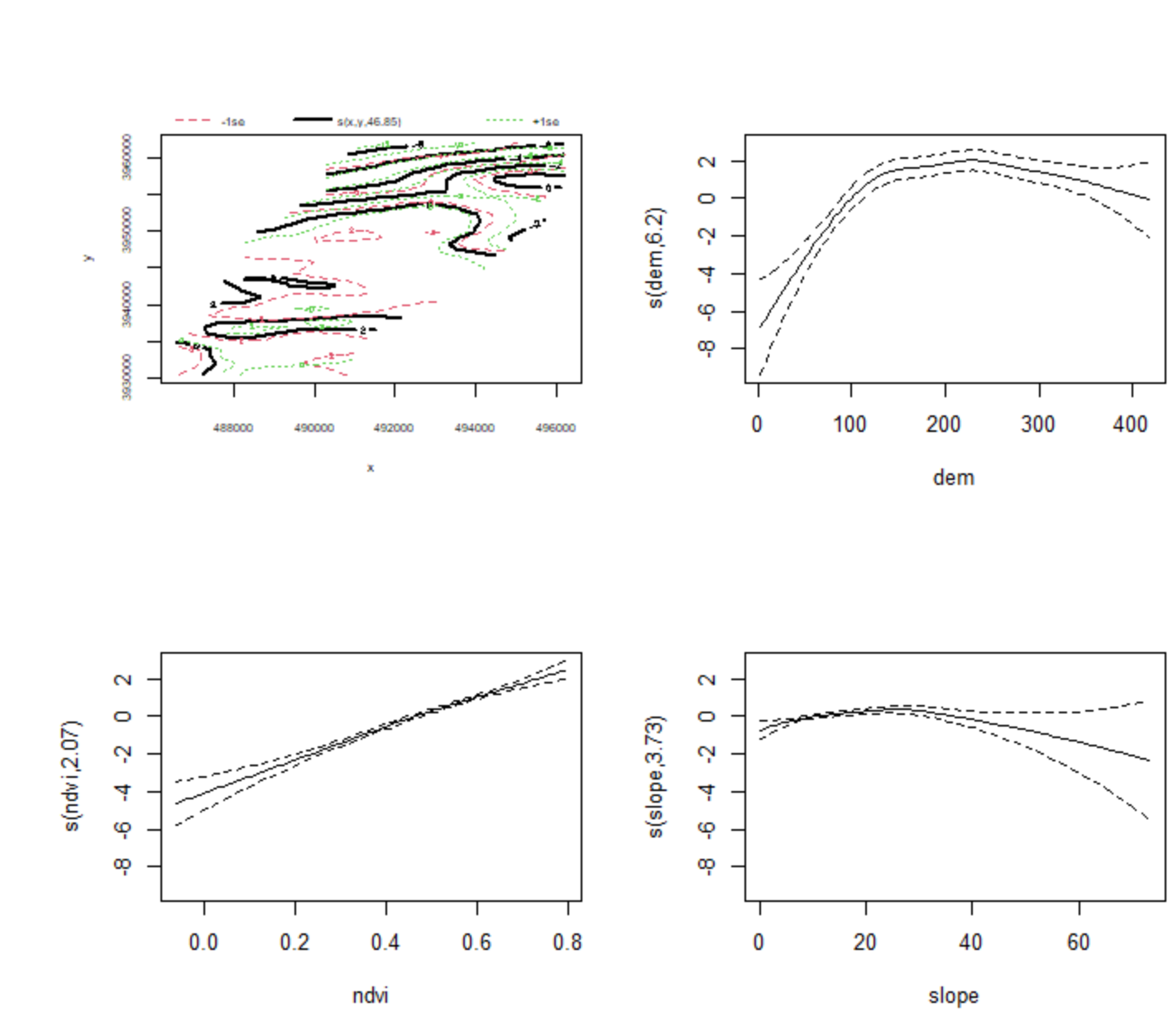

**Figure S4-1:** Partial effects plots of modelled smooth terms from the selected density surface model for Antarctic Prions. Suggests prion density increases at higher NDVI values corresponding with denser vegetation, peak occurrence is between 100 and 300 m elevation, and on moderate slopes.

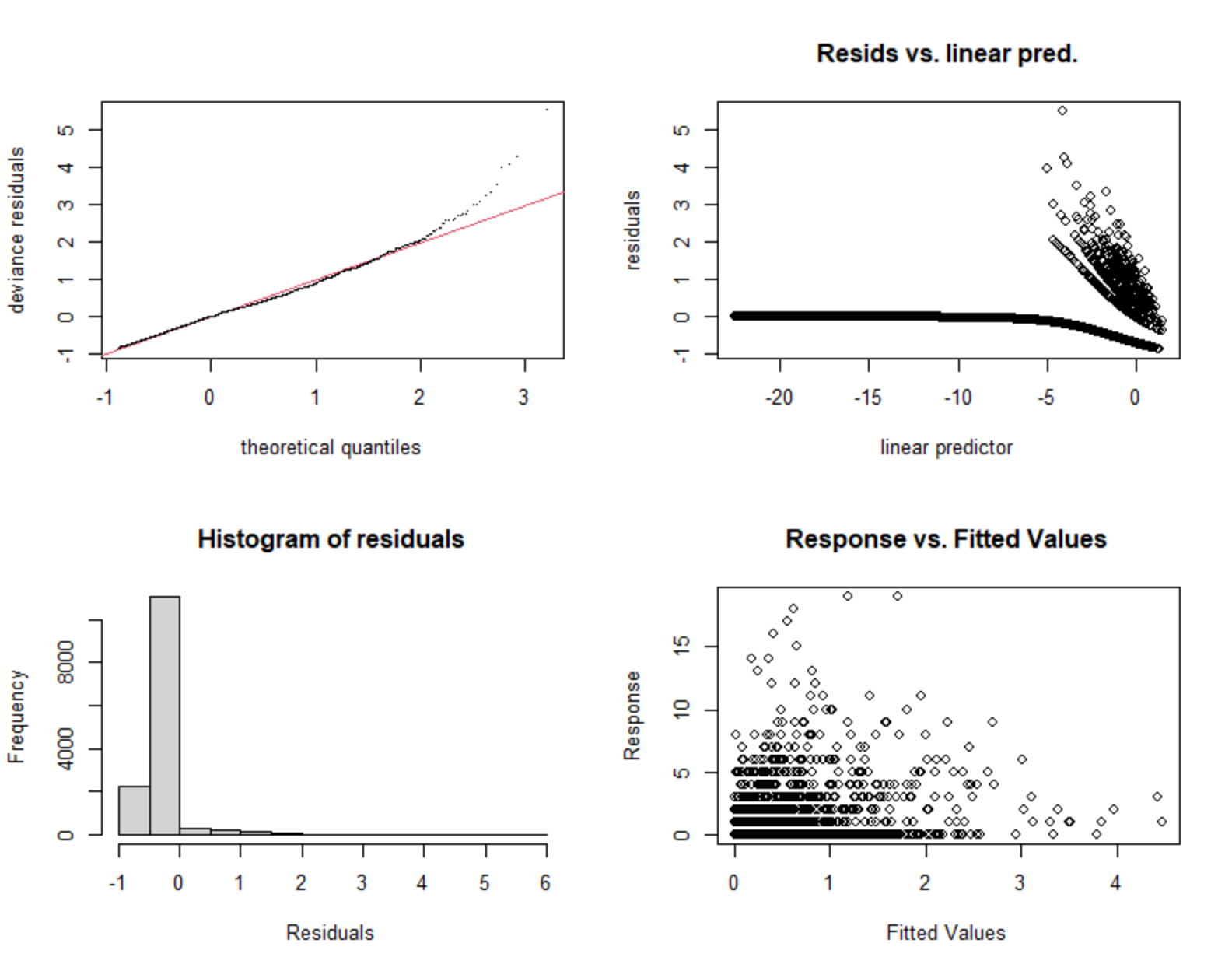

**Figure S4-2:** Diagnostic plots for the selected density surface model for Antarctic Prions. The quantile-quantile plots suggests a reasonable fit between the model residuals and a normal distribution except at high values.

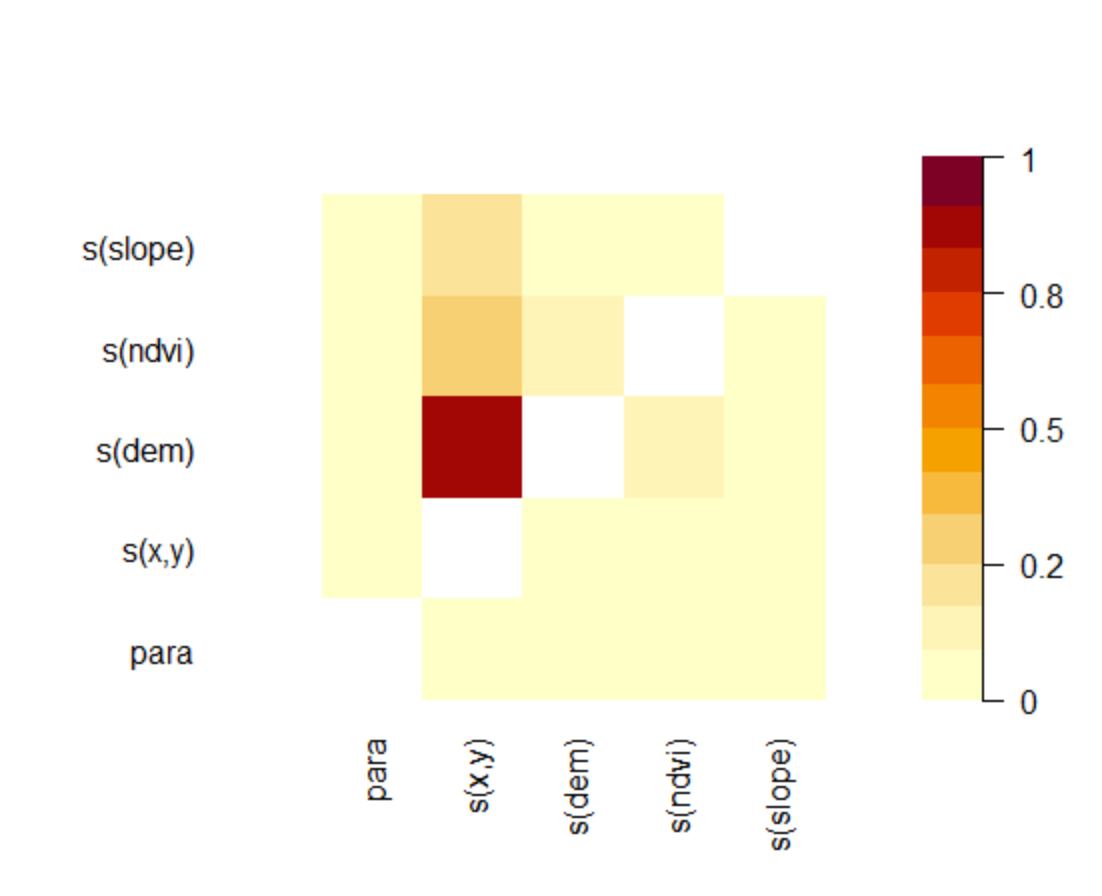

**Figure S4-3:** Concurvity between model terms. The spatial xy terms absorbs much of the variation in other terms, but otherwise they are uncorrelated.

**Table S4-2:** Model selection for White-headed Petrels. Diagnostic plots for global models were compared for three distributions. Then the highest non-significant covariate was dropped until there was no improvement in AIC.

| **model** | **response** | **terms** | **adjusted *R^2^*** | **AIC** | **REML** | **deviance explained** |
| --- | --- | --- | --- | --- | --- | --- |
| dsm_whp_nb | negative binomial | s(x,y), s(dem), s(ndvi), s(ridge), s(slope),  s(wetness), s(wind) | -1.28 | 1389.80 | 711.35 | 84.39% |
| dsm_whp_tw2 | tweedie | s(x,y), s(dem), s(ndvi), s(slope), s(wetness) | 0.35 | 7312.15 | 512.02 | 73.36% |
| dsm_whp_tw5 | tweedie | s(x,y), s(dem), s(ndvi), s(slope), s(wetness) | 0.35 | 7312.15 | 512.02 | 73.36% |
| dsm_whp_tw3 | tweedie | s(x,y), s(dem), s(ndvi), s(slope) | 0.25 | 7563.51 | 641.46 | 71.63% |
| dsm_whp_tw1 | tweedie | s(x,y), s(dem), s(ndvi), s(ridge), s(slope),  s(wetness) | 0.29 | 7568.09 | 639.29 | 72.69% |
| dsm_whp_tw | tweedie | s(x,y), s(dem), s(ndvi), s(ridge), s(slope),  s(wetness), s(wind) | 0.29 | 7570.21 | 640.68 | 72.71% |
| dsm_whp_qp | quasipoisson | s(x,y), s(dem), s(ndvi), s(ridge), s(slope),  s(wetness), s(wind) | 0.70 | NA | -1517.33 | 82.79% |

```
Family: Tweedie(p=1.176)
Link function: log

Formula:
count ~ s(x, y, k = 60) + s(dem, k = 10) + s(ndvi, k = 10) +
    s(slope, k = 10) + s(wetness, k = 10) + offset(off.set)

Parametric coefficients:
            Estimate Std. Error t value Pr(>|t|)
(Intercept) -13.9345     0.7901  -17.64   <2e-16 ***
---
Signif. codes:  0 ‘***’ 0.001 ‘**’ 0.01 ‘*’ 0.05 ‘.’ 0.1 ‘ ’ 1

Approximate significance of smooth terms:
              edf Ref.df      F  p-value
s(x,y)     27.194 34.012  5.363  < 2e-16 ***
s(dem)      5.000  5.928  8.240 7.08e-09 ***
s(ndvi)     4.006  4.969  6.590 4.46e-06 ***
s(slope)    4.160  5.126 12.023 9.66e-12 ***
s(wetness)  2.799  3.602  1.570    0.198
---
Signif. codes:  0 ‘***’ 0.001 ‘**’ 0.01 ‘*’ 0.05 ‘.’ 0.1 ‘ ’ 1

R-sq.(adj) =  0.349   Deviance explained = 73.4%
-REML = 512.02  Scale est. = 2.7608    n = 7140

Method: REML   Optimizer: outer newton
full convergence after 9 iterations.
Gradient range [-7.302387e-05,2.561893e-05]
(score 512.0227 & scale 2.760811).
Hessian positive definite, eigenvalue range [0.5792419,509.1062].
Model rank =  96 / 96

Basis dimension (k) checking results. Low p-value (k-index<1) may
indicate that k is too low, especially if edf is close to k'.

              k'   edf k-index p-value
s(x,y)     59.00 27.19    0.79  <2e-16 ***
s(dem)      9.00  5.00    0.92    0.10 .
s(ndvi)     9.00  4.01    0.95    0.74
s(slope)    9.00  4.16    0.96    0.98
s(wetness)  9.00  2.80    0.91    0.06 .
---
Signif. codes:  0 ‘***’ 0.001 ‘**’ 0.01 ‘*’ 0.05 ‘.’ 0.1 ‘ ’ 1
```

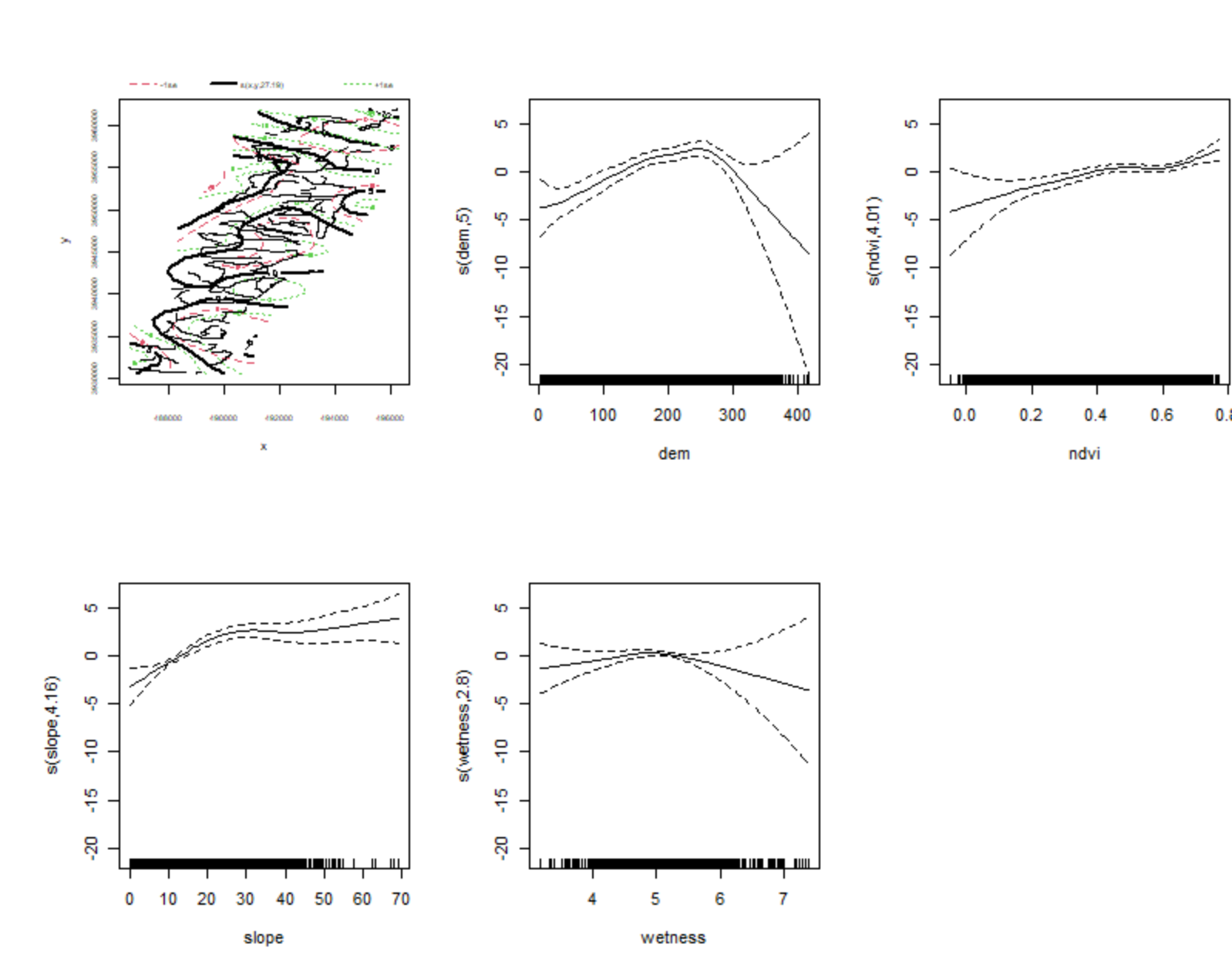

**Figure S4-4:** Partial effects plots of modelled smooth terms from the selected density surface model for White-headed Petrels. Suggests petrel density increases at higher NDVI values corresponding with denser vegetation, peak occurrence is between 100 and 300 m elevation, and on moderate slopes and moderate wetness.

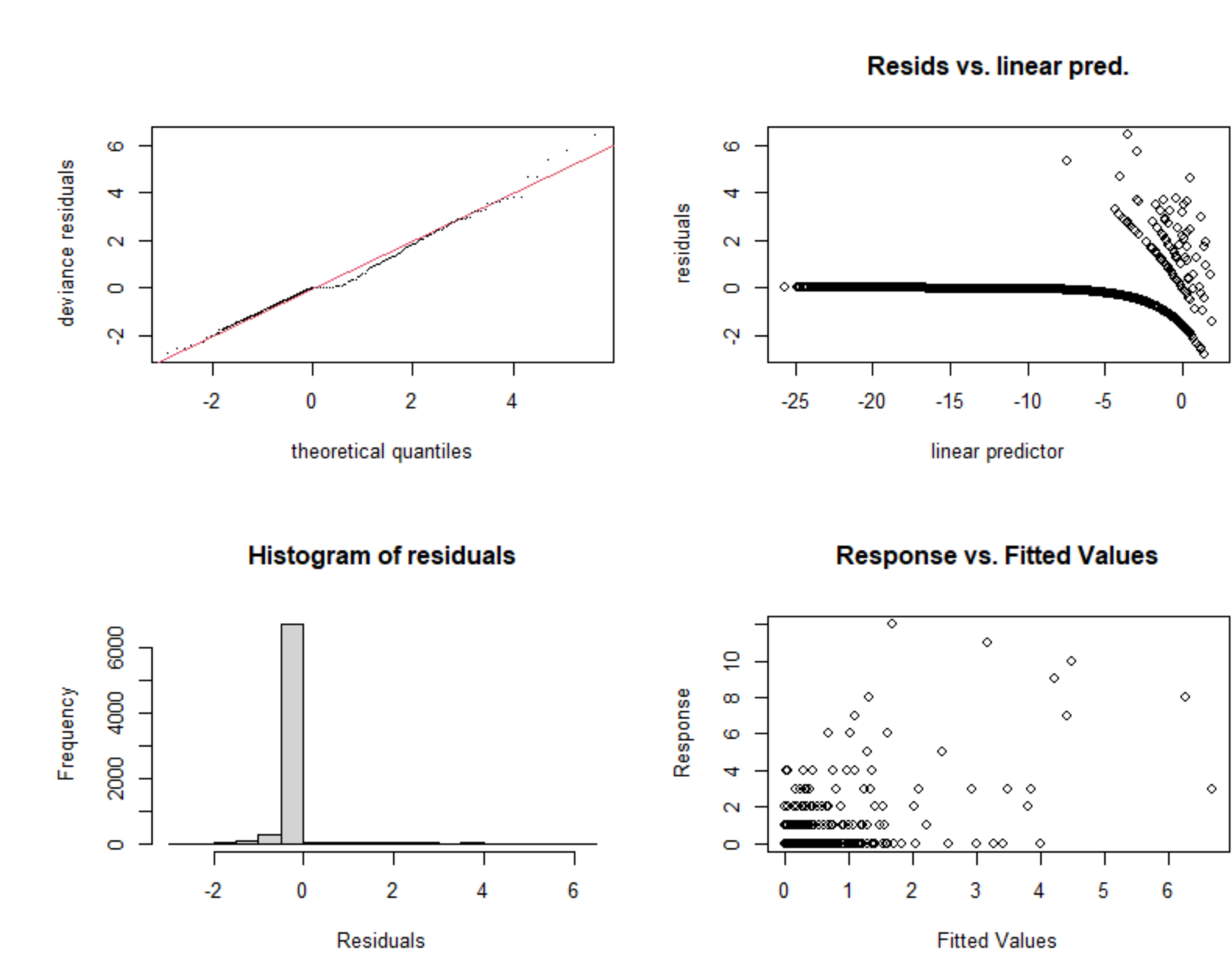

**Figure S4-5:** Diagnostic plots for the selected density surface model for White-headed Petrels. The quantile-quantile plot suggests a reasonable fit between the model residuals and a normal distribution.

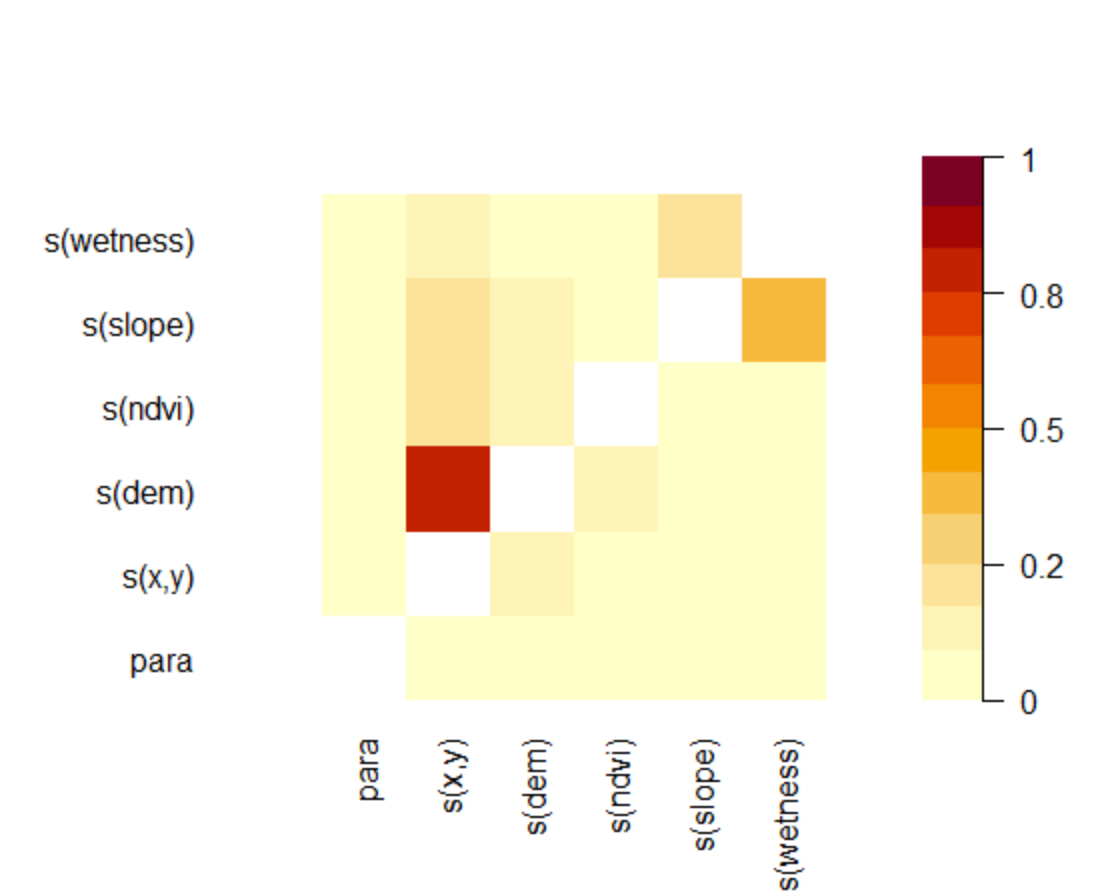

**Figure S4-6:** Concurvity between model terms. The spatial xy terms absorbs much of the variation in other terms, and there is some correlation between slope and wetness.
